# The Optimization of High-Protein Duckweed Cultivation in Eutrophicated Water with Mutualistic Bacteria

**DOI:** 10.1101/2024.02.05.578878

**Authors:** Sunyapat Akkarajeerawat, Thanatkorn Chauvanasmith, Chanan Keatsirisart

**Affiliations:** Bangkok Christian College, Bangkok, Thailand

## Abstract

The exponential increase in the human population is the leading cause of a global food crisis and the rise of humanity’s carbon footprint level. One such approach to resolve these issues is the cultivation of duckweed (*Wolffia globosa*) for consumption in the Northern and Northeastern provinces of Thailand. In particular, duckweed has drawn increasing attention due to its rapid growth rate, adaptability to extreme conditions, and potential as a protein alternative; however, their properties can be greatly enhanced. Our research investigates two optimizations for the cultivation of duckweeds: in the medium’s nitrogen-phosphorus concentration and the use of plant growth-promoting bacteria (PGPB). We hypothesized that using phosphorus-rich water and PGPB would significantly increase duckweed biomass production, while nitrogen-rich water would increase protein content. This was based on evidence from past research showing that aquatic plants have high nitrogen and phosphorus uptake and that PGPB can support the growth of plants by nitrogen fixation. We found that the phosphorus-rich group boosted biomass greatly, while the nitrogen-rich group significantly enhanced their protein value. Furthermore, PGPB yielded the highest increase in biomass, but had an insignificant impact on protein. Based on our results, we recommend the use of nitrogen and phosphorus-rich water and PGPB to optimize duckweed cultivation, addressing the global food crisis and environmental impact.

## INTRODUCTION

The world is currently facing an increase in food shortage due to the exponential population growth and the demand that follow (1). Because of this, finding a promising protein alternative should be the top priority for humanity. Other than proteins from insects, plant-based proteins are also a novel option that are gaining attraction, particularly as the number of vegans are increasing out of concern for animal welfare and their environmental footprint (2, 3). This leads to the production of superfoods, which are foods with high nutritional value due to large amounts of nutrients and bioactive ingredients (4).

Duckweeds (*Wolffia globosa*), categorized as aquatic macrophytes in the Lemnaceae subfamily, are the smallest angiosperms, measuring only 0.4–0.9 mm in length (5). In Thailand, two species of duckweed can be found naturally in silent ponds and marshes, *Wolffia globosa* and *Wolffia arrhiza* (6). Known for their high protein content (45.54% dry weight), duckweeds have become popular superfoods in Thailand (7). Not only do they have a promisingly high level of nutritional value, but they are also one of the fastest-growing plants under ideal environmental conditions; reproducing by vegetative propagation, called fronds (8).

In recent advancements, many different duckweed species are being cultured for their low consumption of water compared to other plant-based protein alternatives, reducing water evaporation rate, and having a water recovery rate exceeding 95% (9). Furthermore, duckweed cultivation can also reduce the production of carbon dioxide, offering a sustainable option that can be cultivated in both polluted and non-polluted waters (9).

Additionally, duckweed cultivation holds a promising approach towards the growing environmental concerns associated with traditional farming practice. The surge in environmental pollution poses a significant challenge for the agricultural community, for example, algae blooms and ocean acidification (10). On one hand, chemical-based fertilizers can enhance the rate of production to supply the growing demand for food. On the other hand, it can cause major health issues and disrupt the ecological balance of the aquatic environment. The accumulation of nitrogen, phosphates, and other fertilizers in water is prone to eutrophication, a destructive process that can cause damage to the agricultural economy (11, 12). However, many studies highlight duckweed in the genus *Lemna*’s ability to efficiently utilize excessive nutrients and adapt to a diverse range of environments, while also remediating pollutants in wastewater (13). Coupled with the use of mutualistic plant growth-promoting bacteria, known to increase the nutrient uptakes of plants via nitrogen fixation and safeguard plants against pathogens, duckweed may be a promising solution to reduce environmental impact while also yielding positive benefits during long-term cultivation (14, 15).

However, with all of this information, little research has been done to address duckweed in the *Wolffia* genus. The purpose of this study is to investigate the effects of nitrogen-phosphorus-rich medium and PGPB on the growth rate and nutritional value of *Wolffia globosa*. We chose to study *Wolffia globosa* due to its abundance and popularity in Thailand, and the lack of research done on the species *Wolffia* in general. We hypothesized that the addition of phosphorus and PGPB would significantly increase duckweed biomass production, while nitrogen would increase protein content. We conducted the research by simulating eutrophication in the growth medium and using commercialized PGPB to create a mutualistic environment. The growth rate aspect of this study was collected from analyzing the change in the average color of the duckweed density, which was heavily adapted from the research of Azetsu & Suetake (16). In this research, we also measured the final duckweed biomass and their protein content.

The hypothesis was supported by our results, which showed that the protein content and biomass production of duckweed significantly increased in the nitrogen-rich group and the phosphorus-rich group, respectively, and that biomass production was significantly boosted in PGPB-enhanced groups. This study could be important to exploring innovative solutions to reduce the challenges associated with sustainable agriculture and food security. Our research can provide an environmentally friendly approach to the optimization of high-protein duckweed cultivation, which could reduce the global food shortage sustainably and cost-effectively.

## RESULTS

We investigated the effects of nitrogen-phosphorus-rich medium and PGPB-enhanced conditions on duckweed biomass production and protein content. We measured the duckweed’s growth rate, net biomass, and nutritional value over 15 days (**Figure 1**).

**Figure 1.**
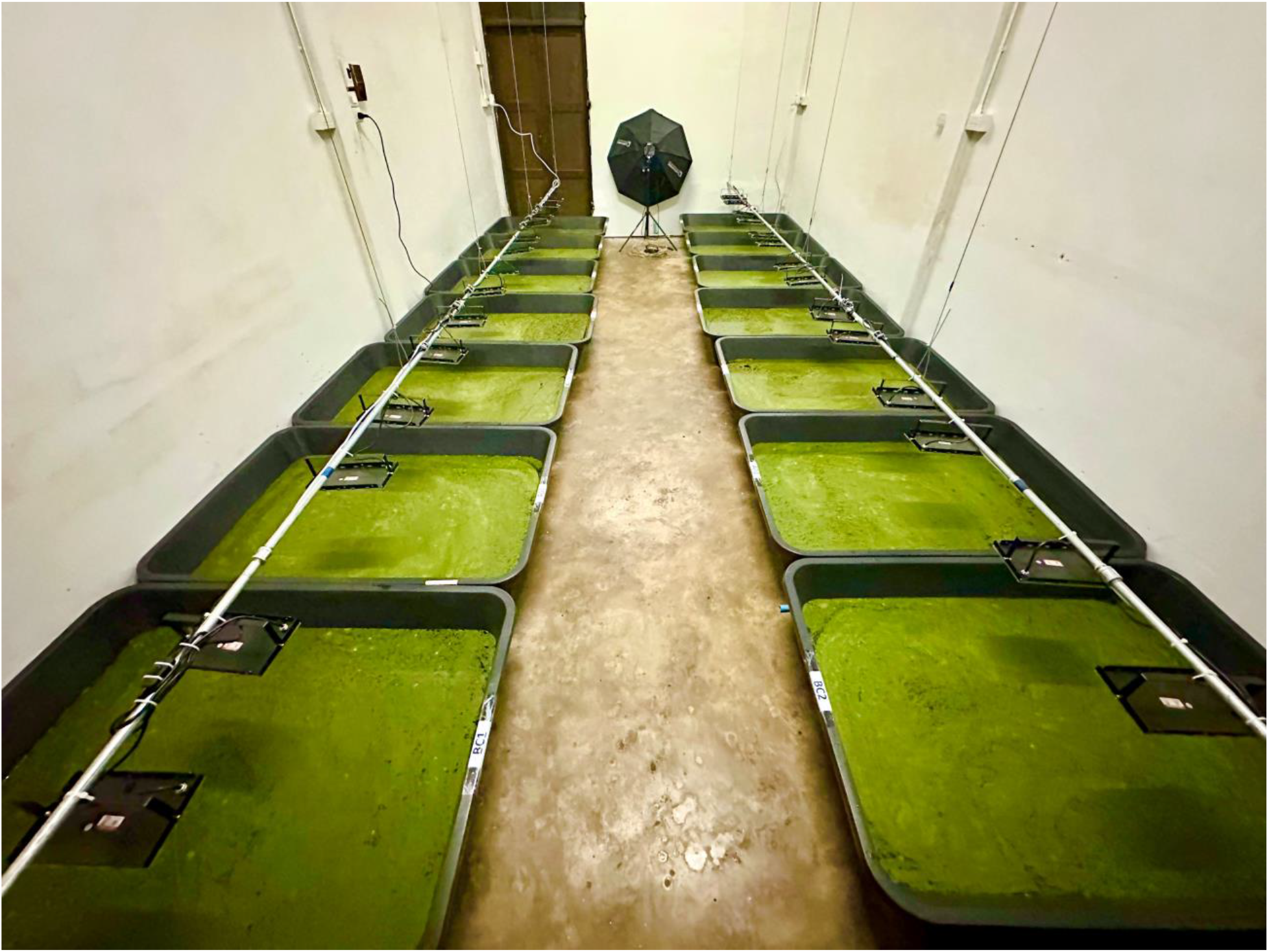
The cultivation of duckweeds (*Wolffia globosa*). *Wolffia globosa* were grown in plastic tubs (93 × 128 × 28 cm) filled with water maintained at a level of approximately 15 cm. inside a controlled room for 15 days.

### Effects of nitrogen-phosphorus-rich media on duckweed

We prepared four different experimental groups: a control group, a nitrogen-rich group, a phosphorus-rich group, and a combined nitrogen and phosphorus-rich group. To effectively measure the growth rate over a large area, we chose to analyze the changes to chroma values in duckweed images (**Figure 2A**). The control group experienced an average change per day of 0.279/day. The experimental samples containing nitrogen and the combined nitrogen and phosphorus display statistically insignificant differences being 0.442/day (*p* = 0.565) and 0.511/day (*p* = 0.329) for the nitrogen-rich group and the combined nitrogen and phosphorus-rich group, respectively. As expected, the phosphorus-rich group significantly elevated the change in chroma value to 0.854/day (*p* = 0.009).

**Figure 2.**
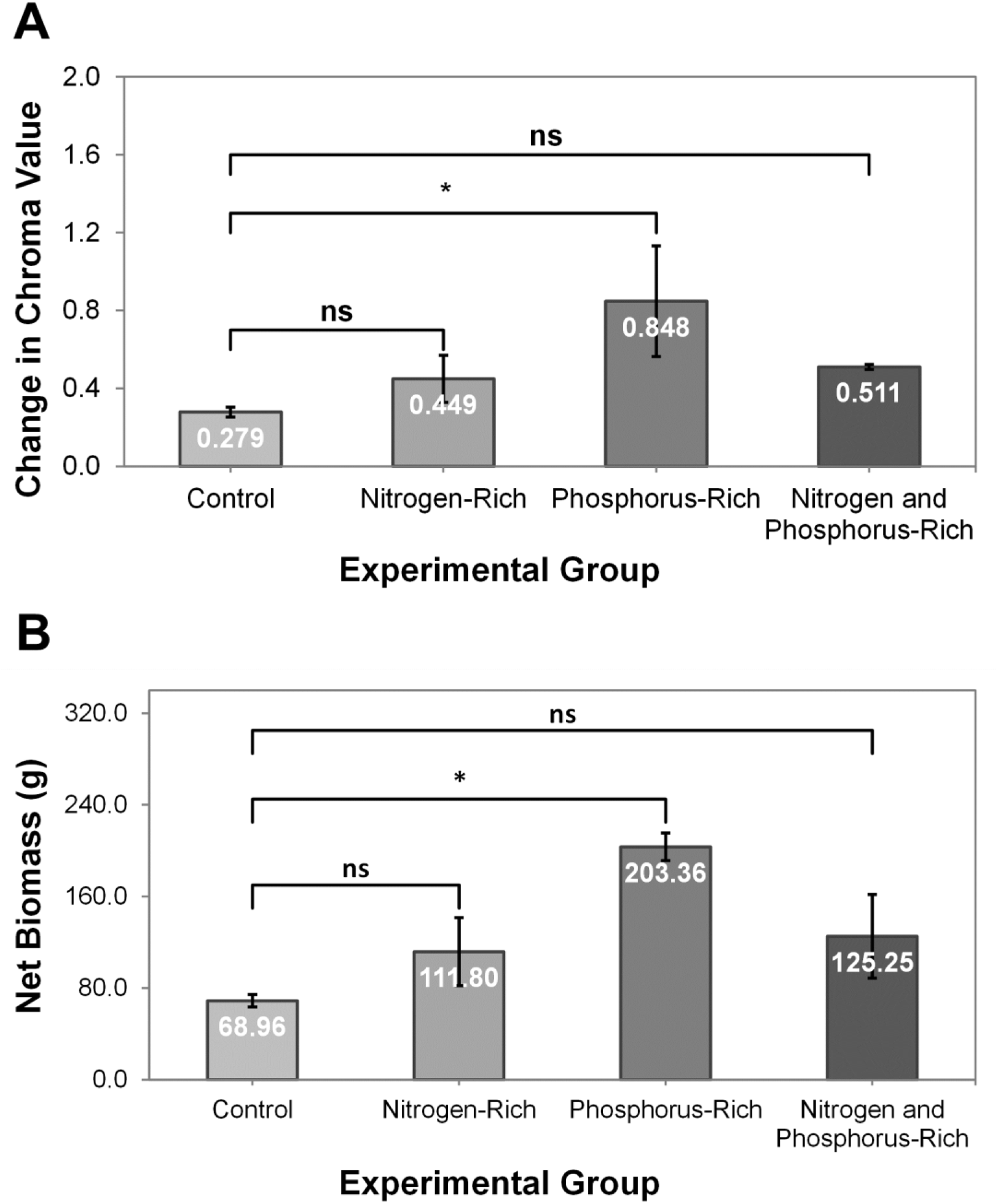
Effects of nitrogen-phosphorus-rich media on duckweed (*Wolffia globosa*) change in chroma value (relative growth rate) and net biomass (n = 3). (A) The change in chroma value of duckweeds (the relative growth rate) cultured in the control group, nitrogen-rich group, phosphorus-rich group, and the combined nitrogen and phosphorus-rich group. (B) The net biomass of duckweeds cultured in the control group, nitrogen-rich group, phosphorus-rich group, and the combined nitrogen and phosphorus-rich group. One-way ANOVA was used to analyze the statistical significance; p > 0.05 (ns) and p < 0.05 (*).

Afterward, we evaluated the net biomass of duckweed after 15 days (**Figure 2B**). The control group showed a net biomass of 68.96 grams. Both nitrogen-rich and the combined nitrogen and phosphorus-rich groups presented an increase in net biomass of 111.80 grams (*p* = 0.217) and 125.25 grams (*p* = 0.085), correspondingly. However, the phosphorus-rich group significantly increased the net biomass to 68.69 grams, which supports our hypothesis (*p* = 0.001).

Next, we analyzed the protein content of duckweed after 15 days. The control group had a protein content of 27.41% (**Figure 3**). The nitrogen-rich group exhibited the most substantial increase in protein content, reaching 52.76% (*p* = 0.001). It was closely followed by the combined nitrogen and phosphorus-rich group with a protein content of 52.50% (*p* = 0.001), while the phosphorus-rich group exhibited a protein content of 41.58% (*p* = 0.029).

**Figure 3.**
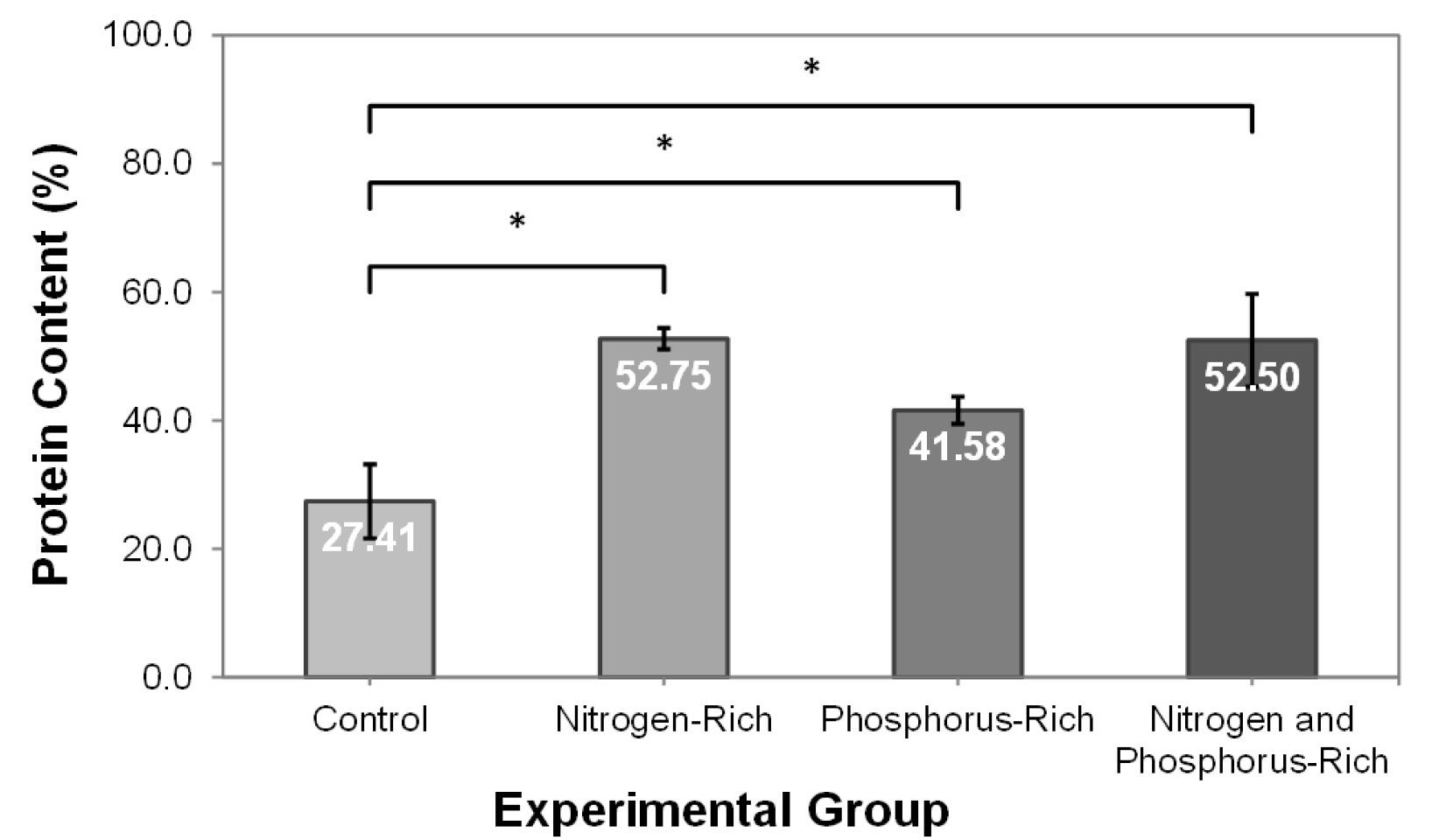
Effects of nitrogen-phosphorus-rich media on duckweed (*Wolffia globosa*) protein content (n = 3). The protein content of duckweeds cultured in the control group, nitrogen-rich group, phosphorus-rich group, and the combined nitrogen and phosphorus-rich group. One-way ANOVA was used to analyze the statistical significance; p > 0.05 (ns) and p < 0.05 (*).

### Effects of PGPB-enhanced conditions on duckweed

We created three experimental groups: a control group, a PGPB-I group consisting of *Azospirillum brasilense* and *Burkholderia vietnamiensis*, and a PGPB-II group comprising *Azospirillum brasilense* and *Gluconacetobacter diazotrophicus*. Upon examining the growth rate of duckweeds, it was found that the control group demonstrated a chroma value change of 0.465/day (**Figure 4A**). In contrast to this, both PGPB-I and PGPB-II groups showed significant changes in their chroma values. Specifically, PGPB-I exhibited an alteration of 1.253/day (*p* < 0.001), while on the other hand, PGBB-II recorded 1.148/day, aligning with our hypothesis (*p* < 0.001).

**Figure 4.**
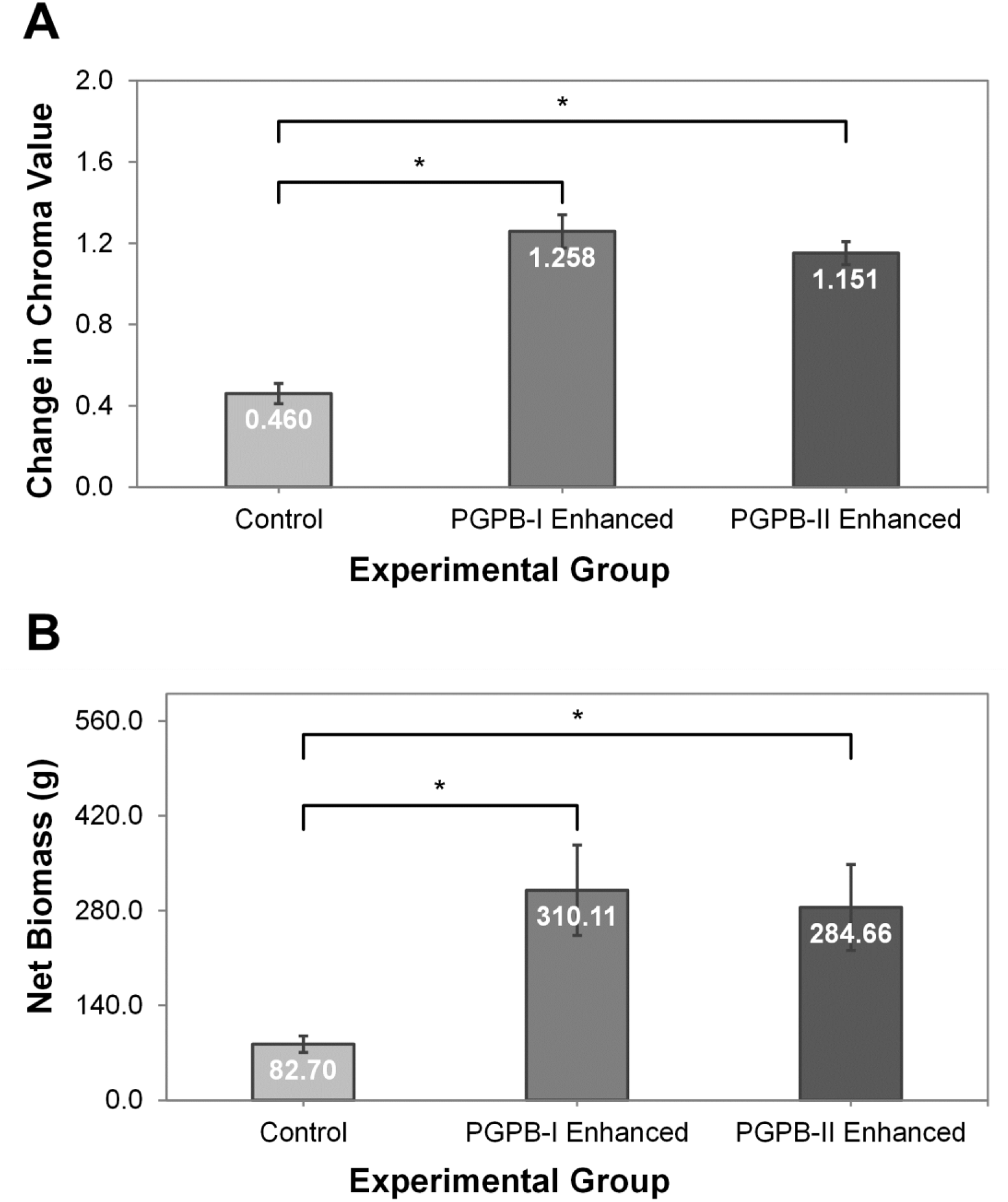
Effects of PGPB-enhanced conditions on duckweed (*Wolffia globosa*) change in chroma value (relative growth rate) and net biomass (n = 3). (A) The change in chroma value of duckweeds (the relative growth rate) cultured in the control group, PGPB-I enhanced group, and the PGPB-II enhanced group. (B) The net biomass of duckweeds cultured in the control group, PGPB-I enhanced group, and the PGPB-II enhanced group. One-way ANOVA was used to analyze the statistical significance; p > 0.05 (ns) and p < 0.05 (*).

Furthermore, with regard to biomass analysis, the control group yielded a net biomass of 82.70 grams (**Figure 4B**). As expected, the PGPB-I group exhibited the highest net biomass output at 310.11 grams (*p* = 0.005), which was comparable to that of the PGPB-II group at a recorded net biomass amounting to 284.66 grams (*p* = 0.009); further supporting our hypothesis.

Finally, we examined the protein content of duckweed (**Figure 5**). The control group displayed a protein percentage of 30.00%. The PGPB-I group exhibited an insignificant increase in protein content reaching 34.67% (*p* = 0.548), while the PGPB-II group achieved higher results with a proportion of proteins amounting to 35.44% (*p* = 0.454).

**Figure 5.**
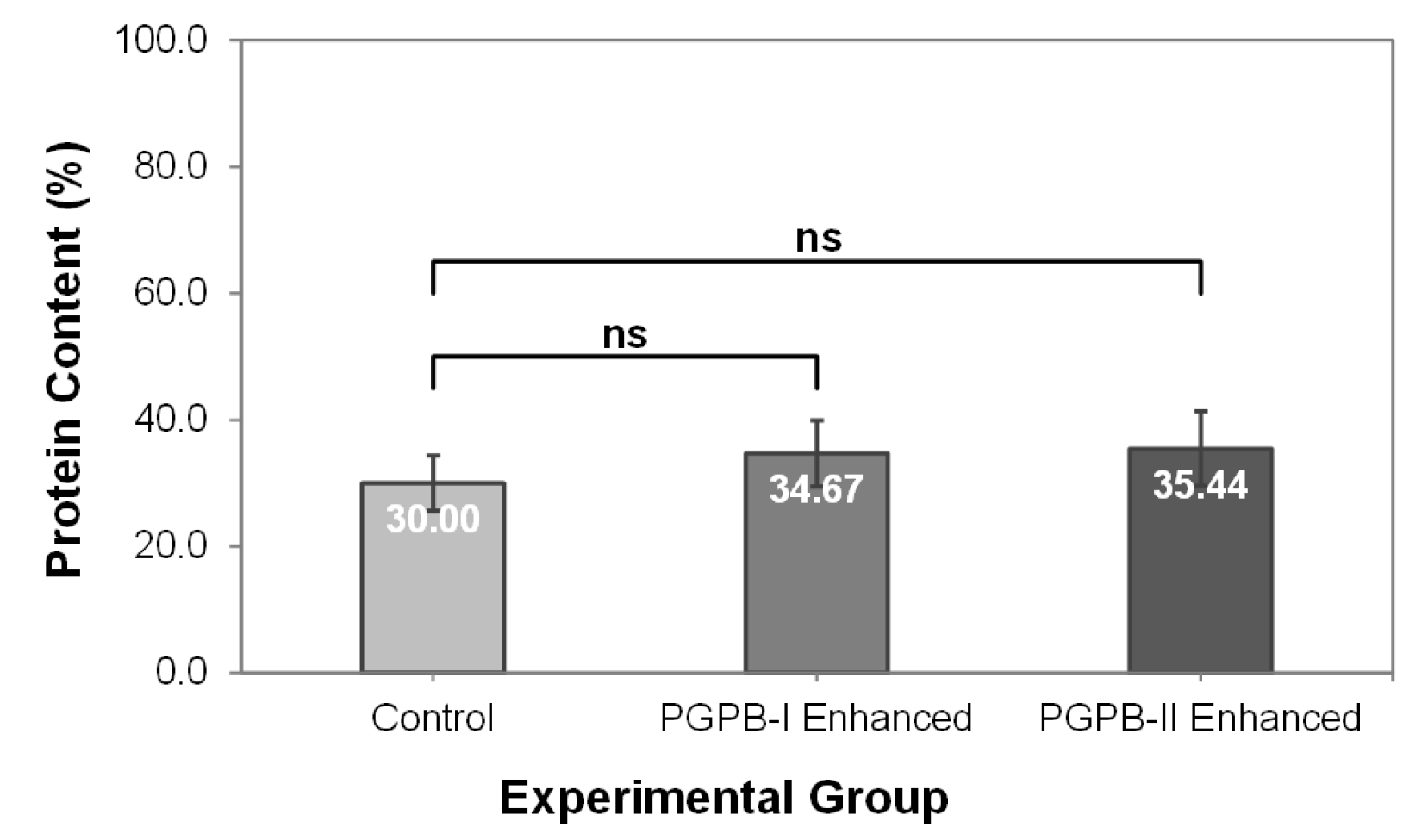
Effects of PGPB-enhanced conditions on duckweed (*Wolffia globosa*) protein content (n = 3). The protein content of duckweeds cultured in the control group, PGPB-I enhanced group, and the PGPB-II enhanced group. One-way ANOVA was used to analyze the statistical significance; p > 0.05 (ns) and p < 0.05 (*).

## DISCUSSION

As the world population is projected to exceed 9.1 billion by 2050 (1), duckweed presents a promising solution for addressing future food shortages. Duckweeds hold nutritional advantages because of their high protein content (approximately 45.54% of dry weight), and considerable amount of carbohydrate and lipids (7). Furthermore, duckweeds contain all nine essential amino acids and have more bioactive ingredients than most grains, such as wheat, corn, or rice (16). In addition, recent research has highlighted *Wolffia globosa*’s functional properties and antimicrobial capabilities within its potential development into functional food ingredients (17).

The present results from this study suggest that duckweed growth is significantly enhanced in a phosphorus-rich medium, as seen through the highest change observed in chroma value at 0.854/day, as compared to the control group which had only 0.279/day (*p* = 0.009). Furthermore, there was an increase in net biomass to 203.36 grams compared to the control group at 68.96 grams (*p* = 0.001). These results align with our initial hypothesis that the addition of phosphorus would significantly enhance duckweed biomass production. However, the data collected from the combined nitrogen and phosphorus group did not exhibit a significant difference compared to the control group. This unexpected outcome could be because of the complex dynamics of duckweed growth, particularly its decline in high-nitrogen mediums. Additionally, due to limitations regarding equipment and time constraints, we were only able to examine one concentration level for each nutrient. Further studies are required to confirm the data as the number of samples tested may have not been enough to fully reveal trends in the larger population.

The protein content of duckweeds grown in nitrogen-rich mediums were significantly increased, with an average of 52.75%, compared to the control group’s 24.41% (*p* = 0.001). Our initial hypothesis was confirmed as the combined nitrogen and phosphorus-rich group and the phosphorus-rich group also demonstrated significant protein content increases. This can be attributed to the biochemical and metabolic functions of nitrogen and phosphorus. Additionally, nitrogen’s impact on chlorophyll and photosynthetic activity may have contributed to this outcome (13).

The effects of PGPB on duckweed growth have also been evaluated. Both PGPB-I (consisting of *Azospirillum brasilense* and *Burkholderia vietnamiensis*) and PGPB-II (consisting of *Azospirillum brasilense* and *Gluconacetobacter diazotrophicus*) significantly enhanced the growth rate of duckweeds, as seen from the high change in chroma value of 1.253/day for the PGPB-I enhanced group (*p* < 0.001), and 1.148/day for the PGPB-II enhanced group (*p* < 0.001), in contrast to the 0.465/day of the control group. Additionally, the net biomass of the duckweeds was also significantly boosted after 15 days, recorded at 310.11 grams (*p* = 0.005) and 284.66 grams (*p* = 0.009) for the PGPB-I and PGPB-II enhanced group, respectively, compared to the control group at 82.70 grams. These results aligned with our hypothesis that the addition of PGPB would significantly enhance duckweed biomass production and support the idea of previous research stating that PGPB can enhance plant nutrient uptake via nitrogen fixation and can safeguard plants against pathogens (15).

While PGPB had a significant effect on improving growth in duckweed, it did not have any notable impact on its protein content. This could be due to limited nitrogen availability in the control medium which prevented PGPBs from nitrogen fixation and resulted in an insignificant yield of proteins. More research is needed with different concentrations of nitrogen to investigate the specific influence of PGPBs on enhancing protein content within duckweeds. Despite some unexpected results during this study, we explored how eutrophicated water combined with the application of PGPBs can optimize both *Wolffia globosa* growth and protein content for the production of superfoods. Moreover, the present study marks the first to investigate the effect of PGPB on *Wolffia globosa* growth and protein content. It is evident that duckweeds have significant potential as a sustainable and cost-effective solution to tackle global food shortages. More comprehensive research in the agricultural industry must be continued for their advancements and improvements so that the hunger crises may one day become obsolete altogether, alongside the current challenges of global warming.

## MATERIALS AND METHODS

### Duckweed Acquisition and Preliminary Preparation

Duckweeds (*Wolffia globosa*) were sourced from a local private farmer in Chonburi, Thailand. Before incubation, the duckweeds were repeatedly washed with filtered water to remove bacteria, algae, and other compounds. The plants were then cultured inside plastic tubs (93 × 128 × 28 cm) filled with water maintained at a level of approximately 15 cm. for 15 days. The parameters were set to 28°C for temperature, light intensity of 12,500 lux, a 12-12 photoperiod, and a pH level of 7. The control cultivation medium consisted of a mixture of 16-16-16 NPK fertilizer obtained from Chemrich, Thailand, and filtered water in a ratio of 1 g: 10 L. All plant cultivations in this study were conducted under the same conditions as above.

### Experimental Groups

After the incubation period, 720 grams of fresh duckweed were divided equally into four groups (each containing 180 grams) to investigate the effect of nitrogen-phosphorus-rich media. The groups included a control group, a nitrogen-rich group (nitrogen concentration of 43.7 mg/L), a phosphorus-rich group (phosphorus concentration of 7.8 mg/L), and the combined nitrogen and phosphorus-rich group (nitrogen and phosphorus concentration of 43.7 mg/L and 7.8 mg/L, respectively). The nutrient concentrations used in this study were based on 53 global wastewater quality datasets (18). The nitrogen and phosphorus used were in the form of soluble urea prills and monocalcium phosphate purchased from Chemrich, Thailand.

For the investigation into PGPB-enhanced conditions, 540 grams of fresh duckweed were divided into three groups (180 grams each). This included a control group, a PGPB-I group containing *Azospirillum brasilense* and *Burkholderia vietnamiensis*, and a PGPB-II group consisting of *Azospirillum brasilense* and *Gluconacetobacter diazotrophicus*. The commercially available PGPBs were purchased from the Department of Agriculture, Thailand.

### Growth Rate Evaluation

To evaluate the growth rate of duckweeds, the plastic tubs were photographed every three days under the same lighting environment and analyzed in Adobe Photoshop 2020 to find the average CIELAB color value of the duckweeds (**Figure 6**). CIELAB, a three-dimensional color space based on human sensitivity, includes three values: *L\** for perceptual lightness and *a\** and *b\** for the four unique colors of human vision (red-green and blue-yellow) (19). The CIELAB values were then used to find the chroma value of the duckweed tubs (the distance of the color from the achromatic axis). As the duckweed tubs are black, an increase in chroma value (the color has a higher intensity and saturation) would indicate duckweed growth. The chroma value was calculated using the following formula.

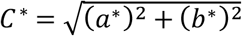

In this equation: *C\** = chroma value, *a\** = red/green coordinate, and *b\** is the yellow/blue coordinate. The change in chroma value (*ΔC\**) was used to evaluate the growth rate of duckweed.

**Figure 6.**
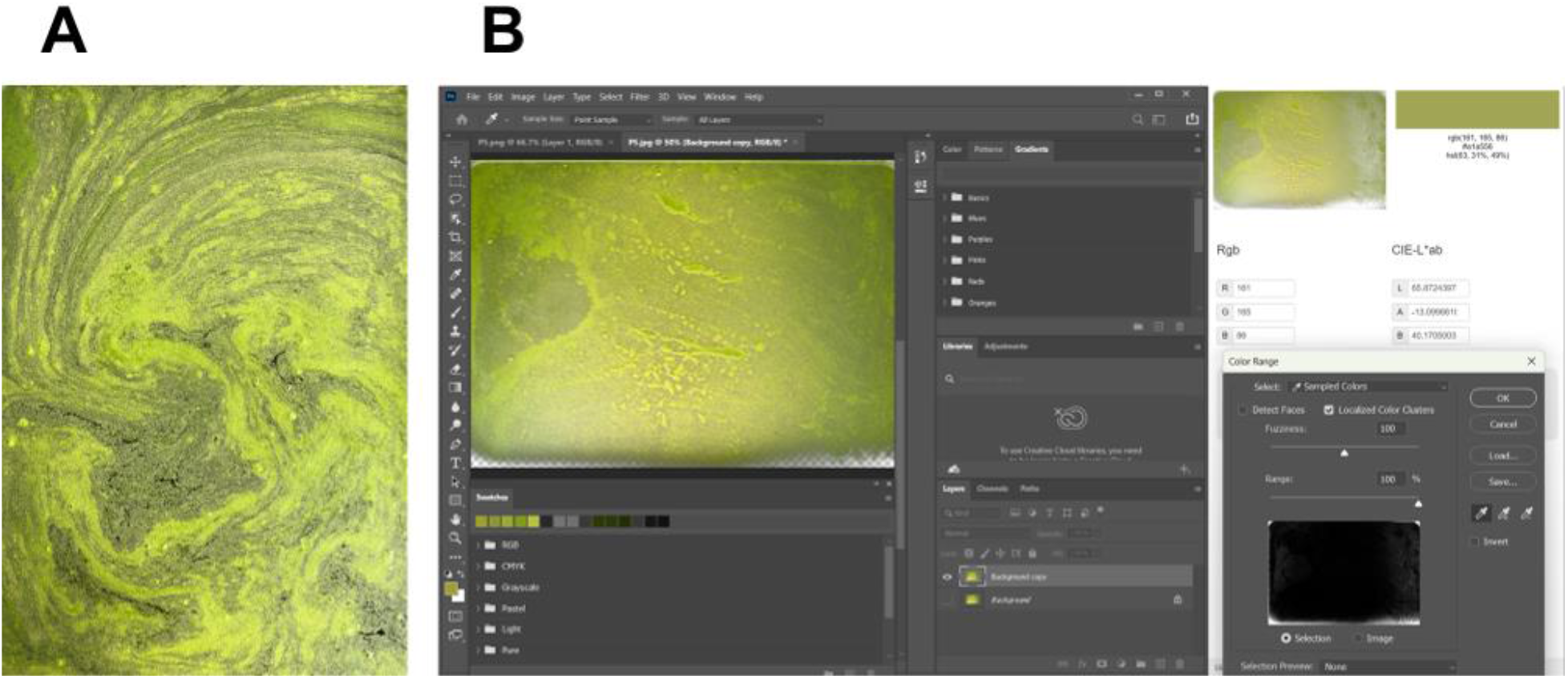
The process of chroma value evaluation in duckweed images. (A) Duckweed tubs were photographed every three days. (B) The duckweed in the images were isolated in Adobe Photoshop 2020. The average CIELAB value of the images were found using an external website.

### Net Biomass Evaluation

To evaluate the net biomass of duckweeds after 15 days, the plants were harvested and dried to remove excess water from the fronds. The final biomasses were weighed using a precision scale (FR-H-1000 digital scale, E-scale). The net biomass was calculated using the following formula.

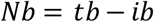

In this equation: *Nb* = net biomass gained after 15 days, *tb* = total biomass of the duckweed obtained after 15 days, and *ib* = initial biomass of the duckweed (which equals 180 grams).

### Protein Content Evaluation

To evaluate the total protein content of duckweeds after 15 days, the biomasses were analyzed by Central Laboratory (Thailand) Co., Ltd. Bangkok, Thailand using an in-house method based on AOAC Official Method 994.12 (2000). The nitrogen content of the samples was determined using the Micro-Kjeldahl method with a conversion factor of 6.25 to calculate crude protein.

### Statistical Analysis

The experiments followed a Completely Randomized Design (CRD) with three replications. IBM SPSS Statistics 29.0 was used to conduct One-Way ANOVA for statistical analysis and Tukey’s Honest Significant Difference (HSD) to discover significant differences (*p* < 0.05) across samples when comparing means.

## ACKNOWLEDGMENTS

We would like to extend our sincere gratitude to the departments and individuals that contributed to the successful completion of this research. We are also grateful to the Science Society of Thailand and Bangkok Christian College for their generous support. Special thanks go to our mentor, Ms. Wanida Bhu-iam, whose invaluable guidance and advice significantly contributed to the research and its statistical analysis.

## Notes

### Competing Interest Statement

The authors have declared no competing interest.

